## Supplemental information for "Denitrification is a community trait with partial pathways dominating across microbial genomes and biomes"

**Contains supplementary tables 1-3; supplementary figures 1-4**

**Supplementary Table 1: Counts and proportion of denitrifier types among different genome sources as a fraction of those encoding *nir* and/or *nosZ***

| denitrifier type | MAG or SCG | isolate | fraction of MAGs or SCGs | fraction of isolates |
|---|---|---|---|---|
| complete | 928 | 1588 | 0.17 | 0.29 |
| NirNor | 1395 | 1512 | 0.25 | 0.27 |
| NirNos | 262 | 154 | 0.05 | 0.03 |
| NirOnly | 1341 | 1252 | 0.24 | 0.23 |
| NorNos | 502 | 356 | 0.09 | 0.06 |
| NosOnly | 1195 | 641 | 0.21 | 0.12 |

**Supplementary table 2: Initial and final selected variables for random forest modeling of gene ratios in marine samples**

| Variable | kept - clade I:II *nosZ* | kept – *nosZ-nir* | minimum | maximum |
|---|---|---|---|---|
| Ammonium (µM) | x | | 0 | 1.93 |
| Sea depth (m) | | x | 20 | 110 |
| Chlorophyll (µg $L^{-1}$) | | | 0.143 | 2.319 |
| Depth (m) | | | 0 | 100 |
| $NO_3^-$+$NO_2^-$ (µM) | | x | 0 | 11.6 |
| $dO_2$ (µM) | | x | 3.614 | 7153.866 |
| Secchi depth (m) | | | 2.5 | 24 |
| Silicate (µM) | x | | 0 | 8.6 |
| Temp (°C) | x | x | 11.34 | 31.13 |
| Total alkalinity (µM) | | x | 2291 | 2367 |

**Supplementary Table 3: Initial and final selected variables for random forest modeling of gene ratios in soils**

| Variable | kept - clade I:II *nosZ* | kept – *nosZ-nir* | minimum | maximum |
|---|---|---|---|---|
| Depth (m) | | | 0 | 0.2 |
| Ammonium (mg $kg^{-1}$) | | | 0 | 87 |
| Conductivity (dS $m^{-1}$) | | | 0.01 | 8.89 |
| Cu (mg $kg^{-1}$) | x | | 0.03 | 32.71 |
| Zn (mg $kg^{-1}$) | x | | 0.02 | 37.65 |
| Elevation (m) | x | | 1 | 1674 |
| SOC (%) | | x | 0.070 | 6.220 |
| pH in $CaCl_2$ | x | | 4.0 | 9.6 |
| Available P (mg $kg^{-1}$) | x | x | 2 | 193 |
| Available K (mg $kg^{-1}$) | x | x | 15 | 1413 |
| Clay (%) | x | | 0.81 | 65.29 |
| $NO_3^-$ (mg $kg^{-1}$) | x | x | 1 | 65 |

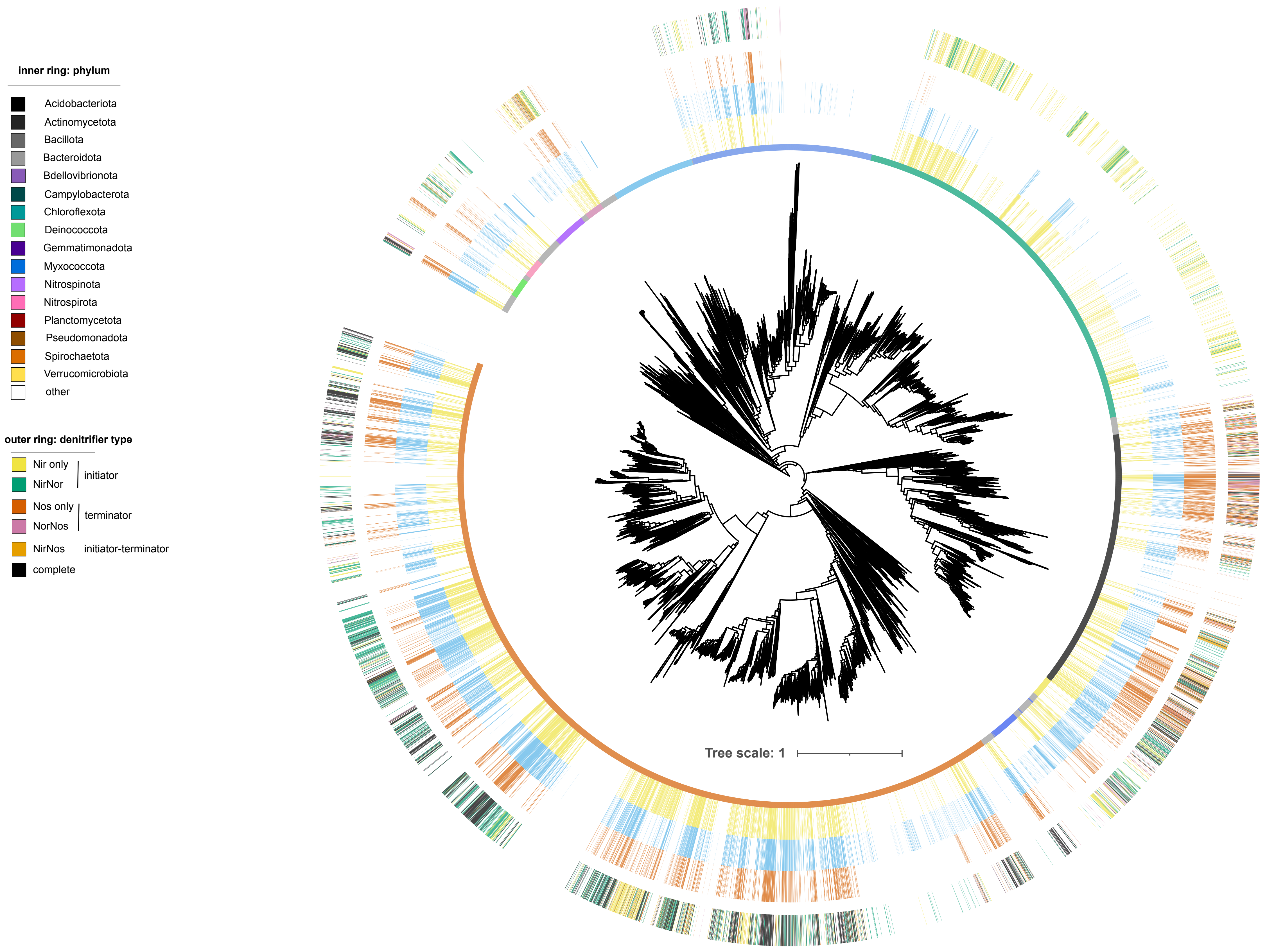

inner ring: phylum
Acidobacteriota
Actinomycetota
Bacillota
Bacteroidota
Bdellovibrionota
Campylobacterota
Chloroflexota
Deinococcota
Gemmatimonadota
Myxococcota
Nitrospinota
Nitrospirota
Planctomycetota
Pseudomonadota
Spirochaetota
Verrucomicrobiota
other
outer ring: denitrifier type
Nir only
NirNor
initiator
Nos only
NorNos
terminator
NirNos
initiator-terminator
complete
Tree scale: 1

**Fig. S1: Phylogeny of representative genomes used in this dataset.** Consecutive rings on tree indicate i. phylum, ii presence of *nirK* and/or *nirS* gene in each genome (yellow), iii presence of nor (BNOR, CNOR; ENOR, GNOR, NNOR, QNOR, SNOR; blue); iv presence of *nosZ* (red) v. denitrifier type. Tree is from GTDB and is based on a reduced set of filtered positions from a concatenated alignment of 120 protein sequences. Scale bar denotes sequence distance based on a JTT+CAT model. Tree was rerooted in Fusobacteria.

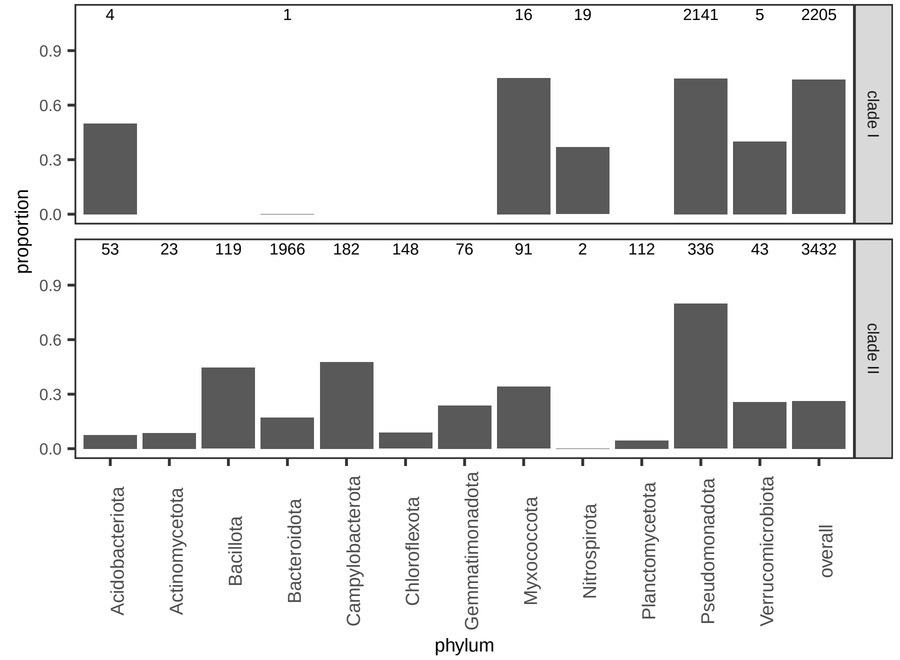

4
1
16
19
2141
5
2205
53
23
119
1966
182
148
76
91
2
112
336
43
3432
0.9
0.6
0.3
0.0
proportion
clade I
clade II
Acidobacteriota
Actinomycetota
Bacillota
Bacteroidota
Campylobacterota
Chloroflexota
Gemmatimonadota
Myxococcota
Nitrospirota
Planctomycetota
Pseudomonadota
Verrucomicrobiota
overall
phylum

**Fig. S2: Differences in prevalence of complete denitrifiers among assemblies encoding clade I vs clade II NosZ.** Bar size denotes proportion of genomes encoding the clade of NosZ that are also complete denitrifiers. Numbers above bars denote number of assemblies. “Overall” denotes average across all genomes encoding that clade of NosZ, and includes data for phyla not plotted individually..

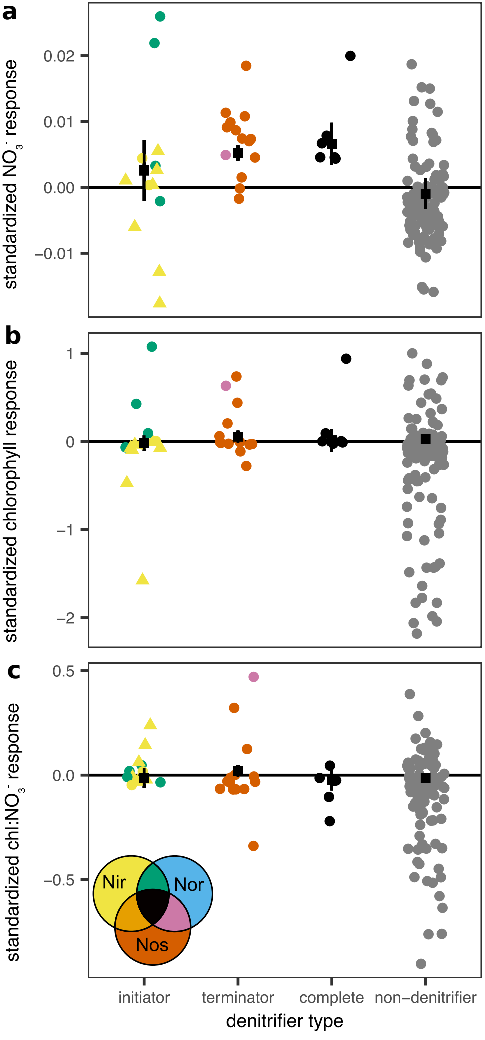
a
b
c
standardized $NO_3^-$ response
0.02
0.01
0.00
−0.01
standardized chlorophyll response
1
0
−1
−2
standardized chl:$NO_3^-$ response
0.5
0.0
−0.5
Nir
Nor
Nos
initiator
terminator
complete
non−denitrifier
denitrifier type

**Fig. S3: Relationship between denitrifier type relative abundance and resource availability in metagenomes derived from marine waters**. **a** nitrate, **b** chlorophyll, and **c** chlorophyll:nitrate ratio. Relative abundance was normalized to the maximum prevalence observed for a given OTU prior to analysis to facilitate comparison between OTUs, and the slope between prevalence and nitrate, chlorophyll or chlorophyll:nitrate ratio was determined for each OTU. Nitrifiers are shown as triangles and all other as circles. Black squares with vertical lines denote weighted mean response and its 95% confidence intervals, respectively, where weighting is proportionate to inverse of the slope standard deviation. The "non-denitrifier" category includes genomes encoding none of the genes and/or just nor. The horizontal lines at zero denote the weighted mean response calculated using the inverse of slope standard error for each value. The two assemblies encoding the initiator-terminator combination Nir-Nos in this dataset are not shown.

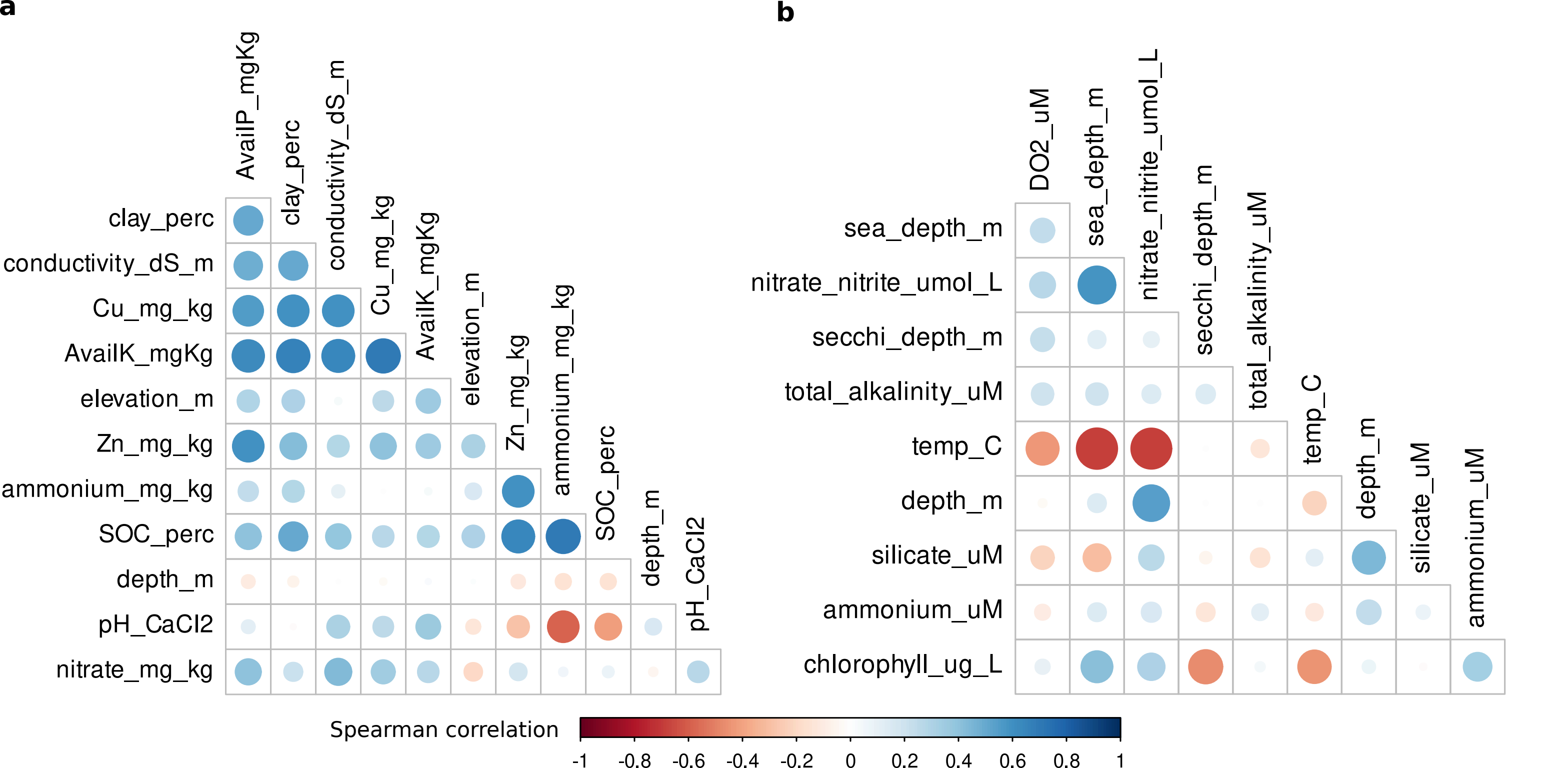
a
AvailP_mgKg
clay_perc
conductivity_dS_m
Cu_mg_kg
AvailK_mgKg
elevation_m
Zn_mg_kg
ammonium_mg_kg
SOC_perc
depth_m
pH_CaCl2
nitrate_mg_kg
b
DO2_uM
sea_depth_m
nitrate_nitrite_umol_L
secchi_depth_m
total_alkalinity_uM
temp_C
depth_m
silicate_uM
ammonium_uM
chlorophyll_ug_L
Spearman correlation
-1
-0.8
-0.6
-0.4
-0.2
0
0.2
0.4
0.6
0.8
1

**Fig. S4:** Spearman correlation between environmental variables used for random forest modeling. a Soils dataset (n=298). b Marine dataset (n=93). Color indicates direction, while intensity is proportional to the correlation coefficients. All correlations are below 0.7. Final variables kept by VSURF and used in random forest modeling can be found in Supplementary Tables 1 and 2.
